# Evi1 is necessary for late activation of delta-notch signaling in sea urchin embryos

**DOI:** 10.1101/2025.04.01.646662

**Authors:** Sanjana Anand, Dr. David McClay

## Abstract

Embryonic mesoderm specification and differentiation rely on intricate gene regulatory networks. In sea urchin embryos, Ecotropic Viral Integration Site 1 (Evi1) is identified as a key transcriptional regulator that integrates Nodal and Delta-Notch signaling pathways to drive mesodermal development. Through in situ hybridization, hybridization chain reactions, and morpholino-mediated knockdowns, Evi1 is required for activating Notch135 expression and for the formation of mesodermal derivatives, including muscle progenitors and coelomic pouches. Evi1 expression is positively regulated by Nodal signaling: it is upregulated when the Nodal antagonist Lefty is knocked down and lost with Nodal depletion or BMP overexpression. Functionally, Evi1 operates upstream of Notch135, restoring its expression following early Delta-Notch activity and enabling a second signaling phase necessary for mesodermal differentiation. Knockdown of Evi1 results in significant loss of Astacin4 and Myo88, markers for mesoderm and muscle progenitors respectively, and disrupts Pax6 expression, impairing coelomic pouch formation, a precursor to the larval rudiment and adult body plan. These results establish Evi1 as a crucial mediator between Nodal and Delta-Notch pathways in mesodermal patterning. Given its conserved role across deuterostomes, further investigation into the direct transcriptional targets of Evi1 and its later-stage developmental functions could shed light on its broader contributions to organogenesis.

## INTRODUCTION

Transcription factors orchestrate gene expression, shaping the developmental trajectories of cells and tissues. Among the many transcription factors studied in sea urchin embryogenesis— a model system with one of the most comprehensively mapped gene regulatory networks— Ecotropic Viral Integration Site 1 (Evi1) remains an enigma. First identified in mammals as a key regulator in leukemogenesis, Evi1 has been linked to oncogenesis (Liang & Wang, 2020), though its role in normal embryonic development is poorly understood. The discovery that Evi1 is expressed in the sea urchin embryo (Rafiq et al., 2014) raises intriguing questions: What does a gene associated with cancer do in an early deuterostome? What role might it play in patterning, differentiation, or signaling? Given that genes implicated in oncogenesis often have critical developmental functions (Liang & Wang, 2020), uncovering Evi1’s role in sea urchin development is not only of fundamental biological interest but may also provide insights into its dysregulation in disease.

This study seeks to define the role of Evi1 in gastrulation, particularly in ventral mesoderm differentiation. Gastrulation is a critical developmental stage during which the three germ layers—ectoderm, mesoderm, and endoderm—are specified through precisely regulated gene expression networks. It was hypothesized that Evi1 interacts with key developmental pathways, including Wnt, Hedgehog, and Delta-Notch signaling, to regulate the formation of critical structures such as the hindgut and coelomic pouches (Peter & Davidson, 2015; McClay, 2011). Using molecular techniques such as in situ hybridizations and hybridization chain reactions (HCR), an examination of Evi1’s expression patterns and functional significance in mesodermal specification shows that Evi1 is a crucial regulator of ventral mesoderm patterning and operates within the Delta-Notch signaling pathway, contributing to muscle and coelomic pouch formation. The implications of our findings position Evi1 at the top of the ventral mesoderm gene regulatory hierarchy, integrating external signals to direct tissue specification.

Mapping Evi1’s function within the Delta-Notch pathway and its interactions with mesodermal and muscle differentiation markers, has advanced the understanding of transcriptional control in sea urchin embryogenesis. More broadly, this work fills a crucial gap in the field and has broader implications for evolutionary developmental biology and human health. Given Evi1’s established role in oncogenesis (Liang & Wang, 2020), understanding its normal developmental function may provide insight into its pathological misregulation.

## METHODS

### Species Collection

Sea urchins (*Lytechinus variegatus*) were harvested from Beaufort, NC, and commercial sources in Florida. Gametes were collected by intracoelomic injection of 0.5 M KCl, and fertilization was carried out in filtered artificial seawater (FASW). Embryos were cultured at 21-23°C in FASW until they reached the desired developmental stages. Embryos were collected at mesenchyme blastula (MB), early gastrula (EG), late gastrula (LG), prism (PM), and pluteus larva (PL) stages for further analysis.

### Embryo Fixation

Embryos were fixed at specified developmental stages in 4% paraformaldehyde in FASW for 30 minutes at room temperature, followed by three washes in phosphate-buffered saline with 0.1% Tween-20 (PBST). Fixed embryos were stored in methanol at −20°C for long-term preservation. Prior to staining, embryos were rehydrated through a graded methanol series into PBST.

### Digoxigenin In Situ Hybridization

Whole-mount in situ hybridization (WMISH) was performed to visualize Evi1 expression using digoxigenin-labeled antisense RNA probes. Probes were synthesized using a DIG RNA labeling kit (Roche Diagnostics) from PCR-amplified templates containing Evi1 sequences. Hybridization was carried out in hybridization buffer (50% formamide, 5X saline-sodium citrate (SSC)), 1 mg/ml yeast tRNA, 100 μg/ml heparin, 0.1% Tween-20) at 60°C overnight. Post-hybridization washes were performed in decreasing concentrations of SSC. Probes were detected using an alkaline phosphatase-conjugated anti-DIG antibody (Roche) and visualized with NBT/BCIP chromogenic substrates. Embryos were imaged using brightfield microscopy.

### Hybridization Chain Reactions

Hybridization chain reactions (HCR) were performed to enhance the detection of Evi1 and associated pathway components, including GataE, Pax6, Hnf1a, and myo88. Embryos were hybridized with DNA probes specific to the target genes, followed by sequential hairpin assembly and fluorescent detection using HCR v3.0 reagents (Molecular Instruments, Inc., Los Angeles, CA, USA). Hybridized embryos were washed thoroughly with SSC buffer and imaged using a fluorescence microscope (Zeiss Axis Imager, München, Germany). Double-staining experiments were conducted using dual probes for Evi1 and Notch pathway markers to confirm co-expression and pathway interactions. Probe specificity was validated by omitting probes as negative controls and comparing expression patterns to prior studies.

### Morpholino Experiments

To investigate Evi1 function, morpholino antisense oligonucleotides (Gene Tools, LLC, Philomath, OR, USA) were designed to block translation. Embryos were microinjected at the one-cell stage with 1 mM Evi1 morpholino or a standard control morpholino (5’-CCTCTTACCTCAGTTACAATTTATA-3’) in 0.2 M KCl. Additional morpholino injections targeted Delta, Nodal, and Lefty to examine their regulatory relationship with Evi1. Injected embryos were cultured at 21°C and collected at different stages for fluorescence imaging and HCR analysis. Control and experimental groups were compared to assess morphological and molecular changes, particularly in ventral mesoderm specification.

### Microscopy

Fluorescence imaging was performed using a Zeiss Axio Imager (Zeiss, München, Germany) equipped with appropriate filters for detecting fluorescence signals from HCR probes and immunolabeled samples. Images were processed using ImageJ software, and fluorescence intensity was quantified to compare expression levels between control and knockdown conditions. Confocal imaging was employed for detailed localization studies using a Zeiss LSM 800 confocal microscope (Zeiss, München, Germany). The experimental approach combined in situ hybridization, HCR, and morpholino-mediated knockdown techniques to elucidate Evi1’s role in sea urchin embryogenesis.

This experimental approach, combining WMISH, HCR, and morpholino-mediated knockdown techniques, enabled precise spatiotemporal mapping of Evi1 expression, its interaction with the Delta-Notch pathway, and its regulatory influence on ventral mesoderm and muscle differentiation.

## RESULTS

### Temporal and Spatial Expression of Evi1 During Embryogenesis

To understand the developmental role of Evi1 in sea urchin embryos, its expression patterns were assessed using HCR. Evi1 expression was observed qualitatively across key developmental stages: MB, EG, MG, LG, PM, and PL. Expression was strongest in the mesoderm and foregut regions during mid- and late-gastrula stages, suggesting a pivotal role in mesoderm and foregut specification (Figure 1).

**Figure 1.**
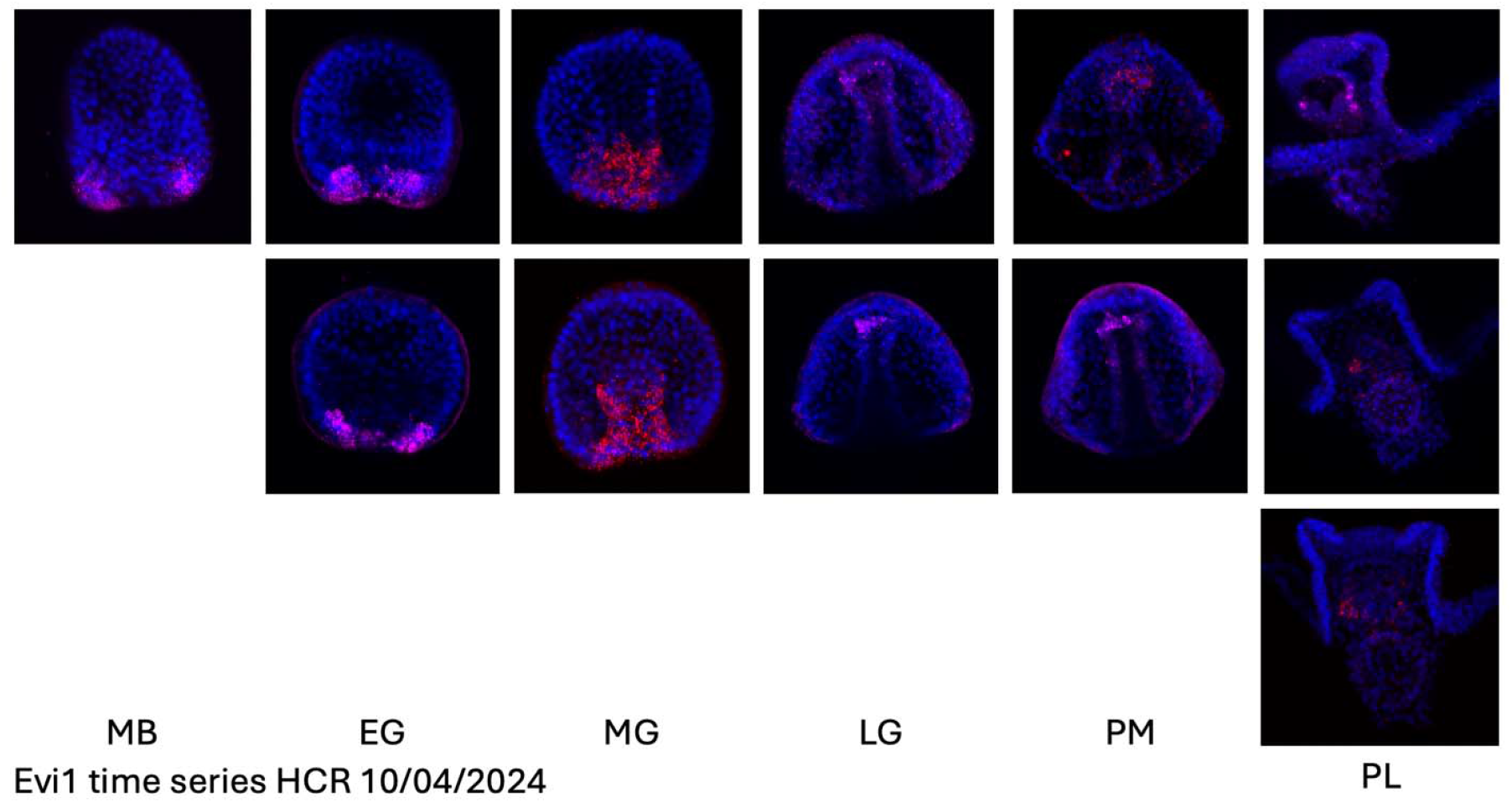
Spatial expression of Evi1 during sea urchin embryogenesis. In situ hybridization images show Evi1 expression across developmental stages: mesenchyme blastula (MB), early gastrula (EG), late gastrula (LG), and prism (PM). Evi1 is highly expressed in the mesoderm region during gastrulation, indicating its role in tissue specification.

### Evi1 and Delta-Notch Pathway Co-Expression

Since Delta induction is necessary for mesoderm specification and that Evi1 is expressed in mesoderm (Figure 1), it is logical to investigate a potential relationship between Evi1 and the Delta-Notch signaling pathway. To explore this, the expression of Notch135, a key receptor in the pathway, was analyzed in Evi1 knockdown embryos. HCR revealed a significant reduction in Notch135 expression following Evi1 knockdown, suggesting that Evi1 positively regulates Delta-Notch signaling (Figure 2).

**Figure 2.**
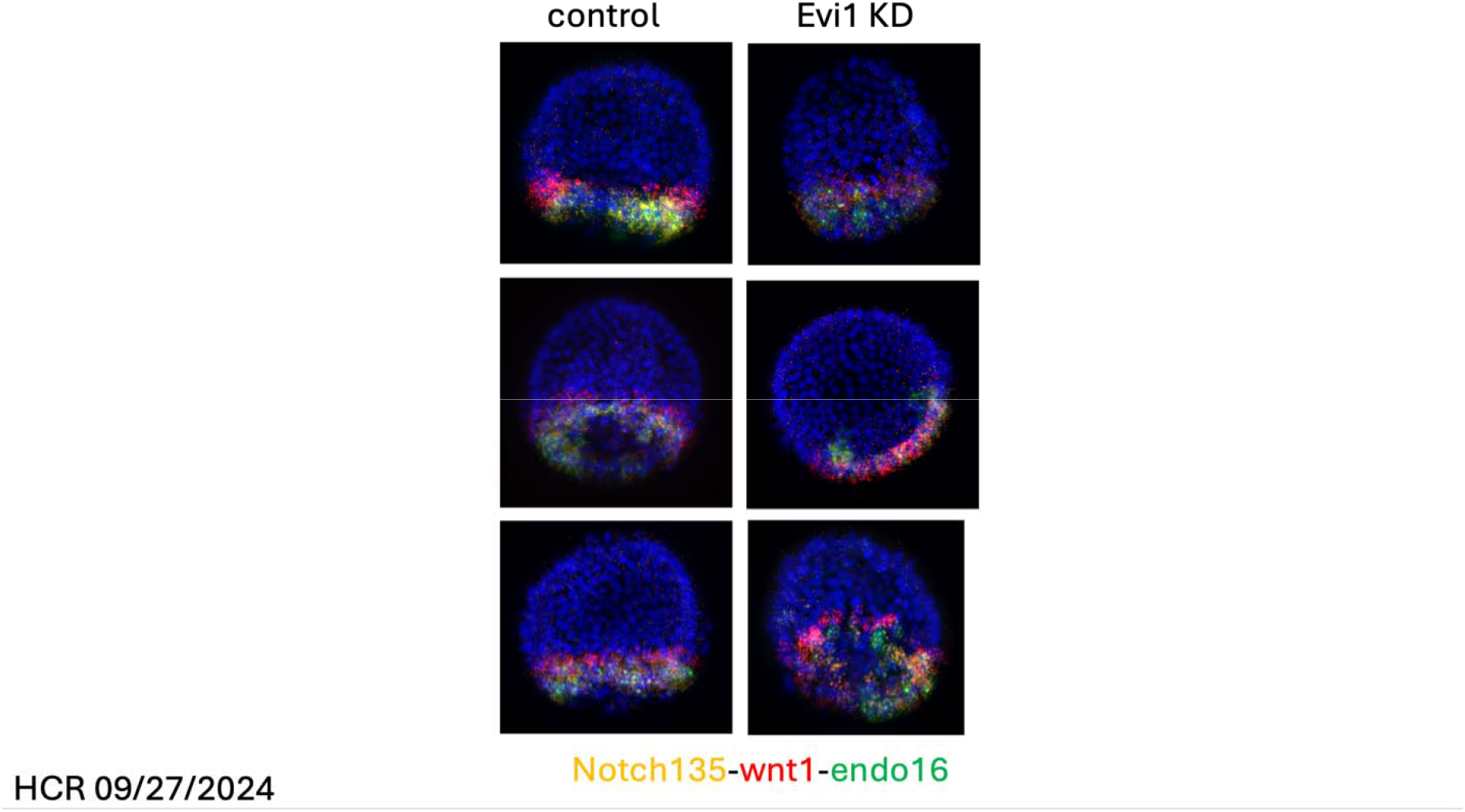
Impact of Evi1 knockdown on mesodermal development. Fluorescence microscopy indicates muted expression of Notch135, a Delta-Notch target gene, in Evi1 knockdown embryos. Wnt1, another signaling component, and Endo16, a marker associated with early mesodermal and endodermal development, were tested, but no significant difference was observed.

To further establish this regulatory link, co-expression studies were conducted, which confirmed that Evi1 and Notch135 exhibit overlapping spatial expression patterns during gastrulation (Figure 3). This spatial correlation implies that Evi1 functions upstream of Notch135, potentially contributing to the presence of Notch.

**Figure 3.**
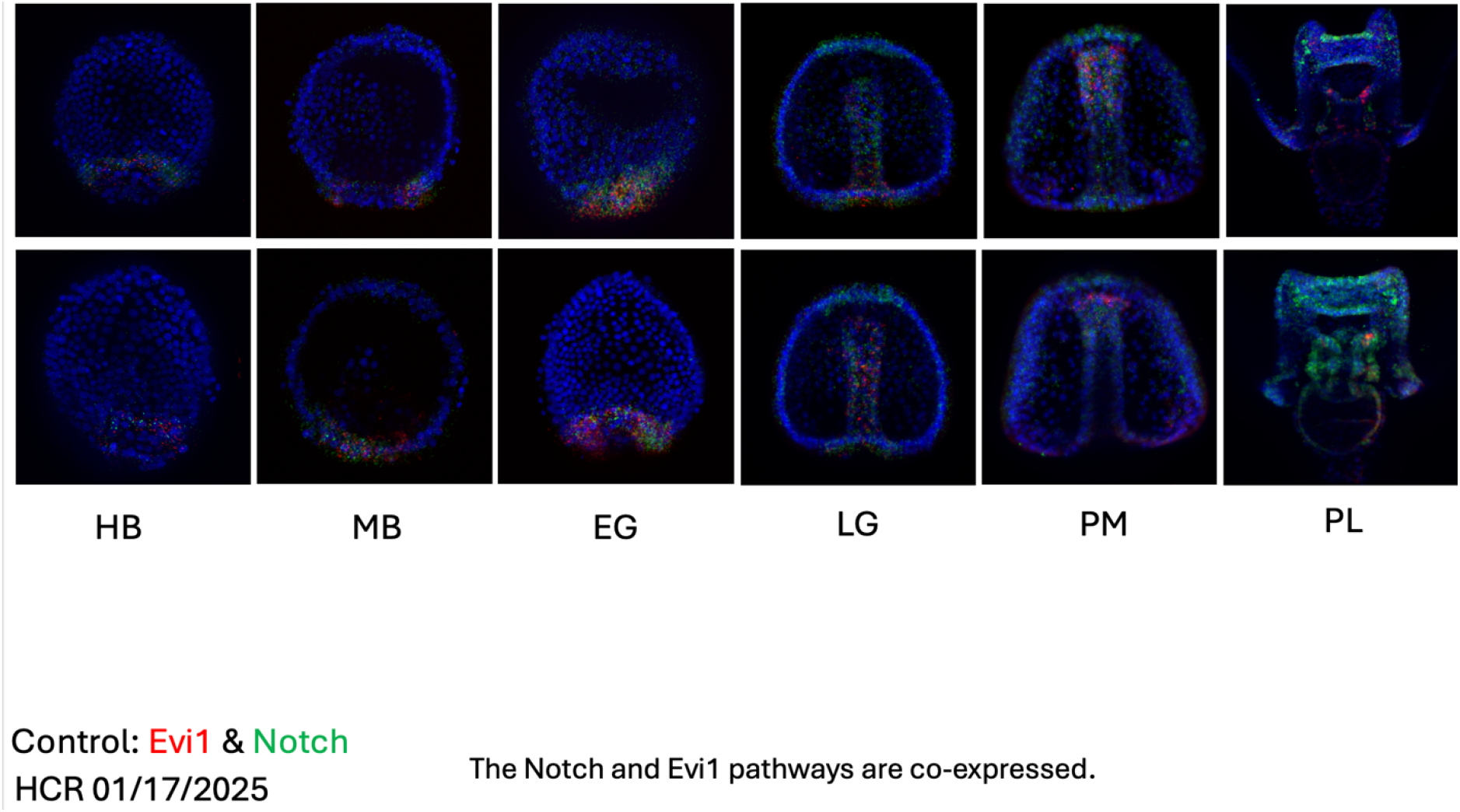
Co-expression of Evi1 and Notch pathway markers. HCR analysis shows overlapping expression of Evi1 and Notch135 during gastrulation.

Given that Notch135 is a critical receptor for Delta-Notch signaling, its loss in Evi1 knockdown embryos indicates that Evi1 regulates the entire Delta-Notch pathway, not just a single component. However, other signaling pathways remain unaffected, as expression of Wnt1 and Endo16, a marker of early mesodermal and endodermal development, was unchanged in Evi1 knockdowns. The Evi1 knockdown of Notch was selective in that two other markers expressed by the same cells, Wnt1 and Endo16, were unaffected by the Evi1 knockdown.

### Evi1 as a Downstream Effector of Nodal Signaling in the Delta-Notch Pathway

The Nodal signaling pathway plays a crucial role in establishing embryonic patterning by directing ventral mesoderm specification and regulating gene expression networks that distinguish ventral from dorsal mesoderm. However, contrary to previous assumptions, Nodal does not directly activate Delta-Notch signaling. Instead, Nodal represses Delta expression in the mesoderm if expressed first, suggesting that Delta-Notch signaling must be activated independently for proper mesodermal differentiation.

To test the relationship between Evi1 and Nodal signaling, Nodal/Lefty/BMP signaling were altered and changes in Evi1 expression were assessed. Knockdown of Lefty, a Nodal antagonist, resulted in an increase in Evi1 expression, consistent with enhanced Nodal activity promoting Evi1 transcription. Conversely, Nodal knockdown or BMP overexpression, which opposes Nodal by outcompeting it for receptor binding and prevents its positive feedback loop, led to an absence of Evi1 expression (Figure 4).

**Figure 4.**
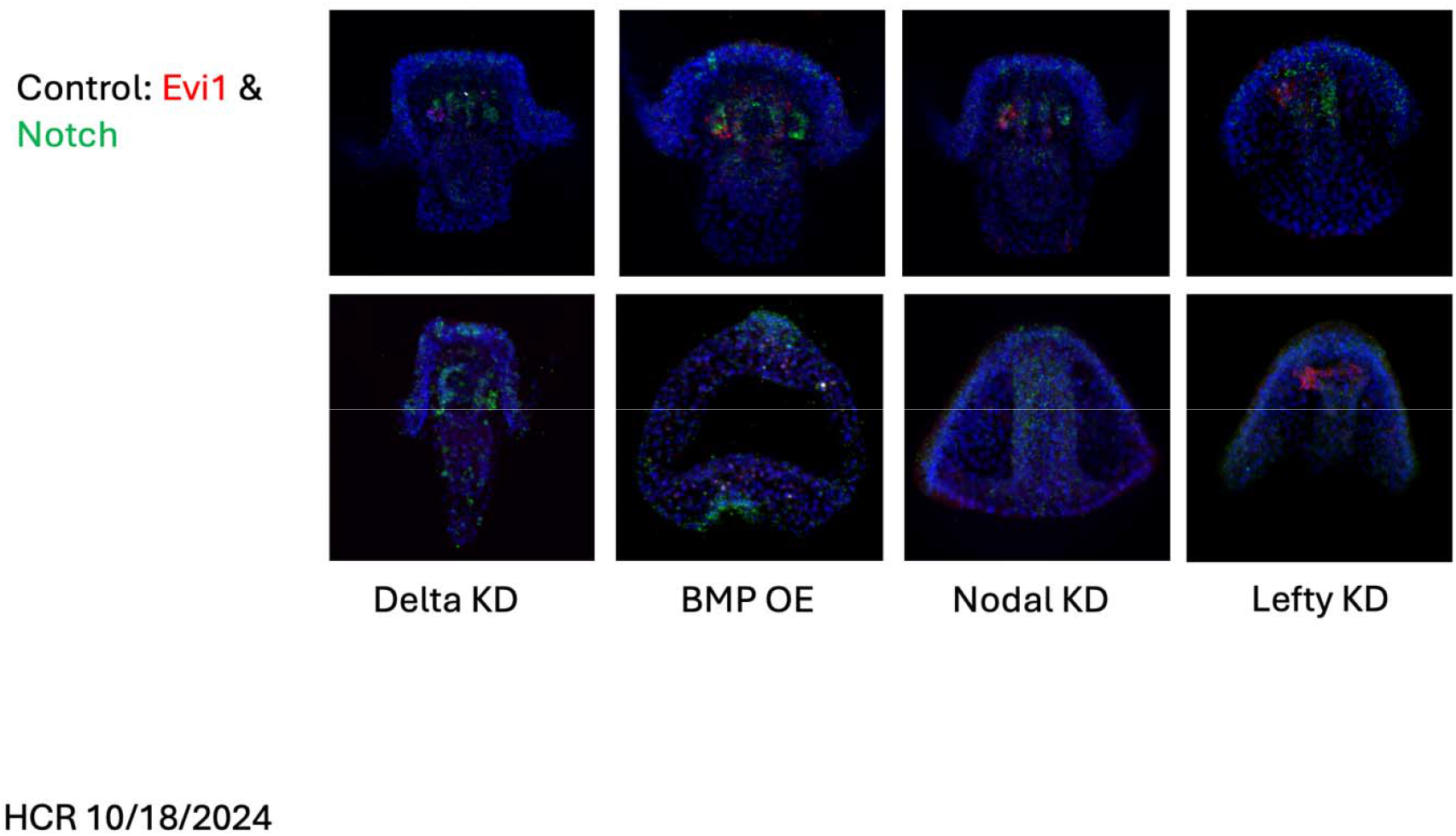
Regulation of Evi1 by upstream signaling pathways. HCR shows co-expression of Evi1 and Notch under control conditions, suggesting pathway interactions. Knockdown of Lefty (causing excess Nodal signaling) leads to overexpression of Evi1, while knockdown of Nodal or overexpression of BMP results in repression of Evi1 expression.

These findings establish that Evi1 is positively regulated by Nodal signaling, positioning it as a key component within the mesodermal gene regulatory network. Normally, Delta-Notch signaling occurs prior to Nodal activation in early mesodermal development, raising the question of how Notch135 is restored in the mesoderm following its initial downregulation. The results provide a possible mechanistic explanation: Evi1 functions downstream of Nodal to re-establish Notch expression, thereby enabling a second phase of Delta-Notch signaling after Nodal activation. This reconciles previous observations that Notch is maternally expressed and present on mesodermal cells prior to Delta induction, after which it is lost from the mesoderm (Sherwood and McClay, 1999). Furthermore, Delta expression in the mesoderm occurs because of early Delta-Notch signaling (Peter & Davidson, 2015), yet Notch itself must be reintroduced to maintain later mesodermal fate decisions. These data suggest that Evi1, regulated by Nodal, is responsible for restoring Notch expression, thereby allowing Delta-Notch signaling to resume following Nodal activation.

Nodal signaling is critical for mesoderm patterning and regulates Delta-Notch signaling to control mesodermal fate decisions. Given its spatial and temporal expression pattern, Evi1 is likely a downstream effector in this network, integrating Nodal and Delta-Notch signals to mediate mesodermal differentiation. However, while Evi1 is expressed near the blastopore during early gastrulation, it is not present in the cells that later contribute to the hindgut. This suggests that Evi1 does not regulate hindgut specification. Whether Evi1 plays a role in foregut development or whether its expression in this region is restricted to mesodermal derivatives remains an open question, as no foregut-specific markers were analyzed in Evi1 knockdown embryos.

### Impact of Evi1 Knockdown on Mesoderm Development

Having established that Nodal is necessary for Evi1 expression, mesoderm specification and muscle differentiation were then investigated through the inspection of Evi1’s downstream function in relation to Nodal. Astacin4, a well-established mesodermal marker, was assessed with mesodermal patterning. Morpholino-mediated knockdown of Evi1 resulted in a significant downregulation of Astacin4, indicating defects in early mesodermal patterning (Figure 5A). These findings strongly suggest that Evi1 is a key downstream effector of Nodal, required for proper mesoderm specification.

**Figure 5.**
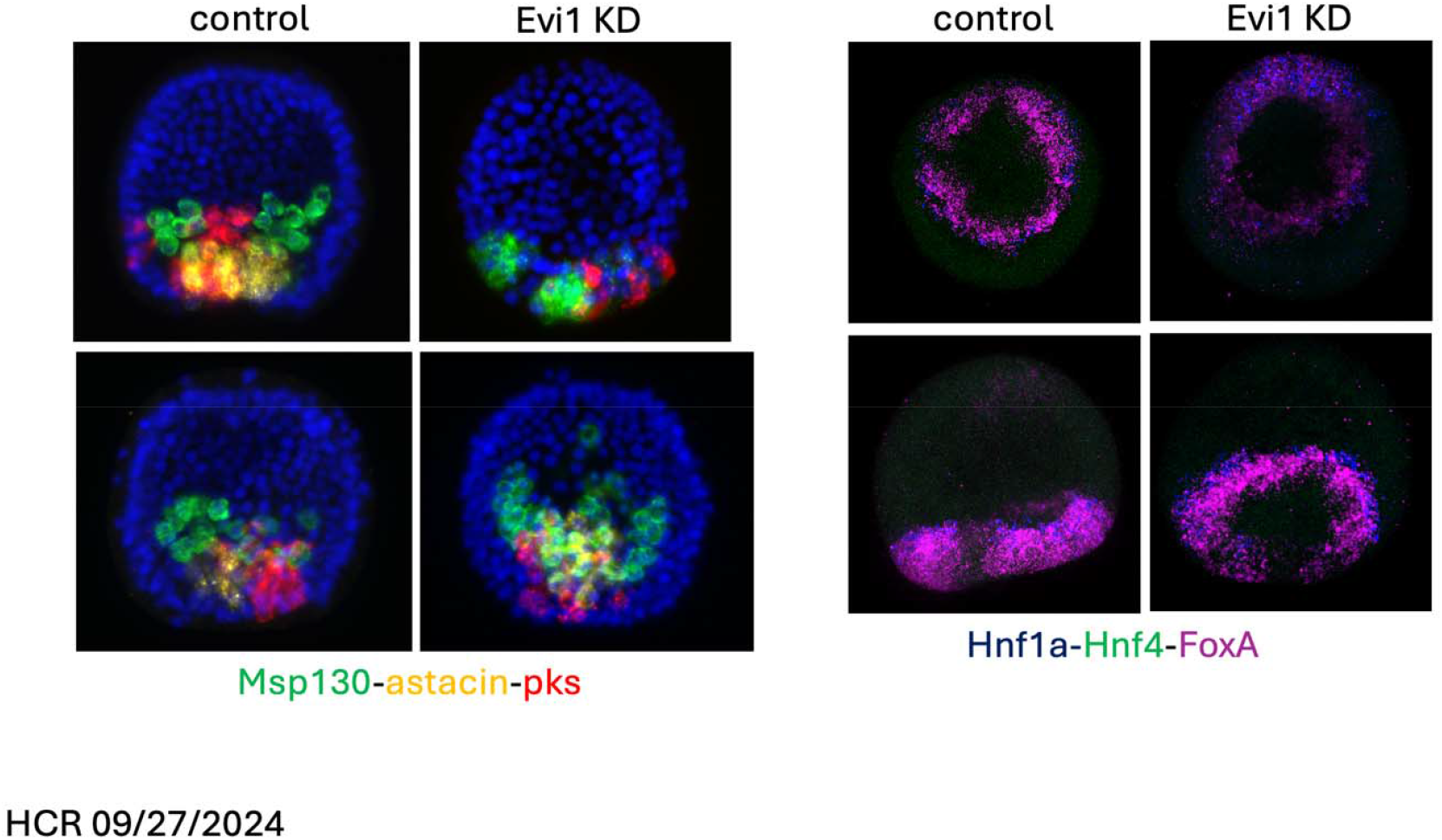
Impact of Evi1 knockdown on mesodermal development. (A) HCR shows reduced expression of Astacin4, a ventral mesoderm marker, in Evi1 knockdown embryos compared to controls. (B) Expression of FoxA, an endodermal marker, remains unaffected, indicating the specificity of Evi1’s regulatory role.

Since muscle is a derivative of mesoderm, Evi1’s role in muscle differentiation was also examined. Myo88, a marker of mesoderm-derived muscle precursors, was completely absent in Evi1 knockdown embryos (Figure 6), suggesting that Evi1 is necessary for the formation of mesodermal muscle progenitors. Myo88 is specifically associated with the development of mesodermal structures adjacent to the coelomic pouches, further linking Evi1 function to mesodermal derivatives. The complete loss of Myo88 expression in Evi1 knockdown embryos suggests that Evi1 is required for the specification of muscle progenitor cells, reinforcing its role in mesoderm differentiation.

**Figure 6.**
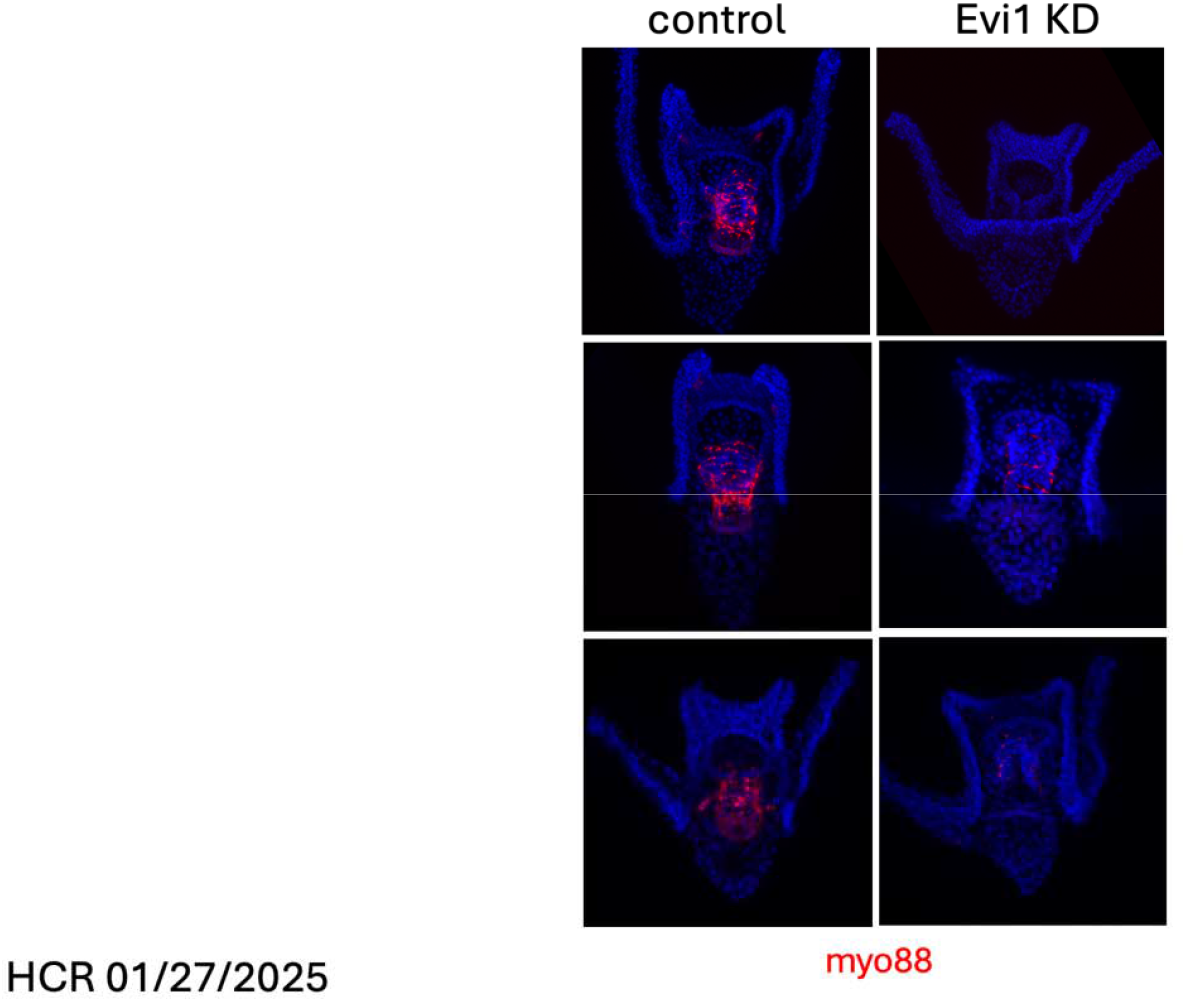
Impact of Evi1 knockdown on mesodermal development. Fluorescence microscopy reveals that Myo88, a muscle-specific gene, is not expressed in Evi1 knockdown embryos, highlighting its role in muscle differentiation.

However, FoxA, an endodermal marker, remained unaffected in Evi1 knockdown embryos (Figure 5B). The use of FoxA as a control was essential to distinguish whether Evi1’s role was specific to mesodermal development or if it broadly affected endodermal tissues. The unchanged FoxA expression supports the conclusion that Evi1 primarily functions in mesodermal specification rather than having a generalized impact on ectodermal or endodermal development.

Taken together, these results establish that Evi1 acts as a critical transcriptional intermediary in the Nodal signaling pathway, regulating both mesodermal specification and muscle differentiation. This supports a Nodal to Evi1 to Astacin4 and Myo88 pathway, in which Nodal signaling activates Evi1, which in turn regulates mesodermal patterning through Astacin4 and muscle development through Myo88. The data also supply a key indication on how Delta-Notch signaling is active after Nodal signaling, by restoring expression of Notch in the mesoderm.

### Evi1’s Role in Coelomic Pouch Specification

Knockdown experiments further revealed that Evi1 is crucial for coelomic pouch development, a mesodermal structure essential for the formation of the adult sea urchin body plan. The left coelomic pouch gives rise to the larval rudiment, which later differentiates into the adult structures during metamorphosis. To investigate Evi1’s role in this process, we analyzed the expression of Pax6, a well-established marker of coelomic pouch formation. Evi1 knockdown embryos exhibited significantly reduced Pax6 expression (Figure 7), suggesting defective specification of coelomic pouch progenitor cells.

**Figure 7.**
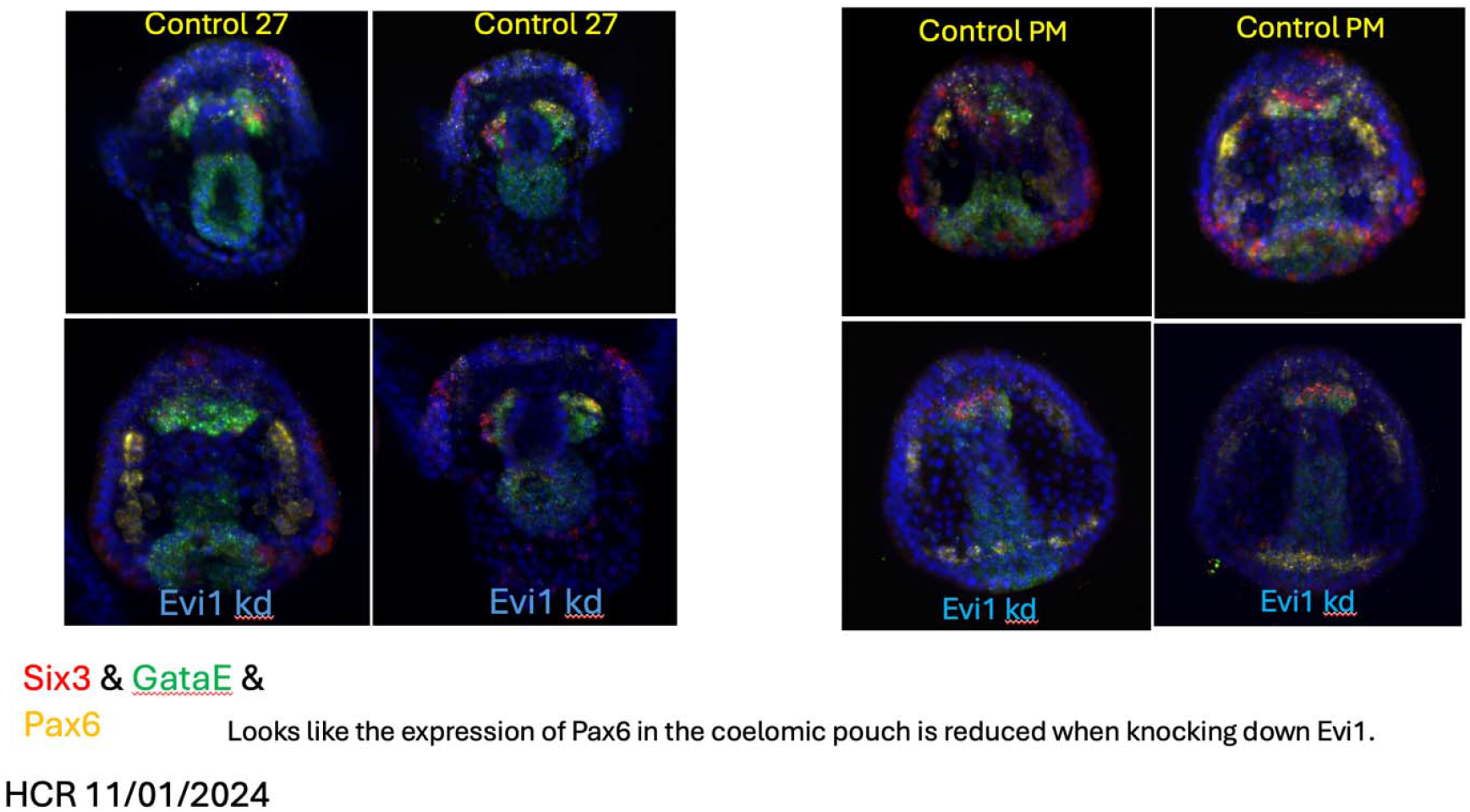
Impact of Evi1 knockdown on mesodermal development. Fluorescence microscopy indicates muted expression of Pax6, a coelomic pouch marker, in Evi1 knockdown embryos.

**Figure 8.**
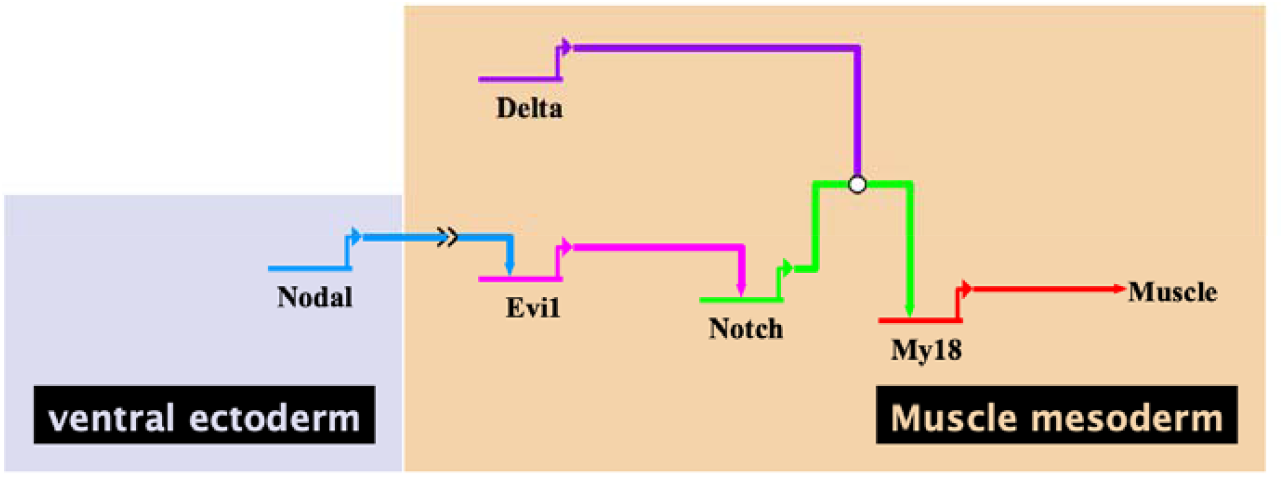
Gene regulatory network illustrating the role of Evi1 in muscle specification. Each horizontal line represents a gene and its associated 5’ regulatory apparatus (enhancer and promoter). Arrows from one gene to another gene’s enhancer depict transcriptional regulation— activation or repression. Nodal signaling, initiated in the ventral ectoderm, activates the transcription factor Smad2/3, which in turn activates Evi1 expression in the mesoderm. Evi1 then activates Notch, restoring its expression following the earlier downregulation of maternally supplied Notch. Delta provides the Notch signaling input necessary to activate Myo18, which ultimately drives muscle differentiation. Perturbation experiments targeting each of the upstream components resulted in the loss of downstream gene expression, validating the inferred regulatory relationships and providing the empirical basis for this gene regulatory model.

Given that Evi1 expression was previously observed in the coelomic pouch region (Figure 4), particularly in the left pouch that later forms the rudiment, this reduction in Pax6 expression reinforces the role of Evi1 as a regulator of coelomic pouch formation. The loss of Pax6 expression in Evi1 knockdowns indicates that Evi1 is necessary for establishing the progenitor cell population that contributes to rudiment formation and, ultimately, adult body structure.

## DISCUSSION

### Evolutionary and Developmental Significance of Evi1

The function of Evi1 in mesodermal specification appears to be conserved within deuterostomes, though its presence and role in non-deuterostome lineages remain unclear. While Evi1 is well established as a regulator of hematopoiesis and leukemogenesis in vertebrates (Liang & Wang, 2020), its function in early mesoderm differentiation has been less extensively characterized. These findings suggest that Evi1 plays a critical role in sea urchin mesodermal development, positioning it as a potential regulator of mesodermal fate in other deuterostomes. Studies in zebrafish and mammals indicate that Evi1 interacts with key developmental pathways during early mesodermal patterning (Goyama et al., 2008; Lenkov et al., 2011). However, evidence for its function in protostomes or non-bilaterian animals such as cnidarians is lacking, and further comparative studies will be necessary to determine whether Evi1’s regulatory role is specific to deuterostomes or has deeper evolutionary origins.

### Evi1 Functions Upstream of the Delta-Notch Pathway

Delta-Notch signaling is essential for early mesodermal fate decisions, and through a combination of in situ hybridization, HCR, and morpholino-mediated knockdown experiments, our findings provide new insight into how Notch expression is re-established in mesodermal cells. Previous research demonstrated that Notch is maternally supplied and initially present as a protein in mesodermal cells, where it facilitates mesoderm specification rather than directly driving Delta-Notch signaling (Sherwood & McClay, 1999). Following this early signaling event, Notch is lost from mesodermal cells, as shown by antibody staining. The mechanism by which Notch expression is later restored to allow a second wave of Delta-Notch signaling remained unclear. The study demonstrates that Evi1 is required for re-establishing Notch expression in mesodermal cells, thereby enabling the continuation of Delta-Notch signaling after Nodal activation.

The reduction in Notch expression in Evi1 knockdown embryos supports the hypothesis that Evi1 functions upstream of Notch, restoring its expression after it was lost following the initial maternal supply. This restoration is crucial for enabling a second phase of Delta-Notch signaling necessary for mesodermal differentiation. Additionally, co-expression studies confirm spatial overlap between Evi1 and Notch during gastrulation, reinforcing this regulatory relationship. To ensure that the observed effects were specific to Evi1 and not due to broad disruptions in mesodermal gene regulation, we assessed the expression of Wnt1 and Endo16, two markers associated with early mesodermal and endodermal development. Their expression remained unaffected in Nodal knockdown embryos, confirming that the absence of Evi1 is directly attributable to the loss of Nodal signaling rather than widespread perturbation of gene expression. These findings expand on previous research showing that Notch signaling is required for mesodermal differentiation (Croce & McClay, 2010) by demonstrating that Evi1 plays a critical role in re-establishing this pathway through transcriptional regulation.

### Evi1 is Essential for Mesoderm Specification

Evi1’s expression pattern during gastrulation highlights its mesoderm-specific function, as it is absent from ectodermal and endodermal lineages. This is further supported by the observation that FoxA, an endodermal marker, remains unaffected in Evi1 knockdown embryos, confirming that Evi1 does not broadly influence endodermal fate. Instead, Astacin4, a well-characterized mesodermal marker, was significantly downregulated in Evi1 knockdown embryos, indicating defects in early mesodermal patterning. While genomic studies have previously reported the presence of Evi1 in deuterostomes (Rafiq et al., 2014), no prior research has identified its function in mesodermal fate decisions. This study is the first to establish that Evi1 is required for mesoderm specification and differentiation, revealing its critical role in the gene regulatory networks governing early embryonic development.

### Evi1 is Required for Muscle Differentiation and Coelomic Pouch Formation

The loss of Myo88 expression in Evi1 knockdown embryos suggests that Evi1 is necessary for the formation of mesoderm-derived muscle progenitors. Given that Myo88 is specifically expressed in developing muscle cells, its absence in knockdown embryos provides strong evidence that Evi1 functions as a determinant of muscle fate. Additionally, the observed reduction in Pax6 expression in the coelomic pouch further supports Evi1’s role in mesodermal tissue specification, as coelomic pouch progenitors give rise to essential adult structures. These results align with genomic studies that have reported Evi1 expression in the sea urchin embryo (Rafiq et al., 2014); however, no prior research has identified its function in mesodermal differentiation.

This study demonstrates that Evi1 plays a crucial role in coelomic pouch development, as evidenced by the significant reduction in Pax6 expression following its knockdown. Evi1 expression was detected in the left coelomic pouch, which later forms the larval rudiment, indicating that Evi1 is essential for establishing progenitor populations that contribute to the adult body plan. The observed loss of Pax6 expression in Evi1 knockdown embryos suggests that Evi1 is required for the proper specification of coelomic pouch progenitors, reinforcing its role in mesodermal differentiation. While coelomic compartments originate from mesoderm, their contribution to specific structures in different phyla varies. The role of Evi1 in coelomic pouch development highlights its importance in mesodermal patterning, ensuring the correct specification of progenitor cells that contribute to adult sea urchin structures.

## Conclusion

This research provides the first mechanistic evidence that Evi1 functions at the intersection of Nodal and Delta-Notch signaling in mesoderm specification. These findings resolve a longstanding question regarding how Notch signaling is re-established following its initial downregulation, positioning Evi1 as a key regulator of mesodermal fate, muscle differentiation, and coelomic pouch formation. The discovery of Evi1’s role in sea urchin embryogenesis has broad implications for both developmental biology and evolution, as its conserved function in deuterostomes suggests regulatory parallels in vertebrate systems.

While Evi1 is established as a transcriptional regulator linking Nodal and Delta-Notch signaling, several questions remain regarding its broader developmental role. Future studies should investigate whether Evi1 directly influences foregut development, as no foregut-specific markers were analyzed in Evi1 knockdown embryos. Although FoxA is expressed throughout the gut, including the foregut, its unaltered expression in Evi1 knockdowns does not clarify whether Evi1 directly regulates foregut fate or is restricted to mesodermal patterning. Transcriptomic approaches could help identify direct transcriptional targets of Evi1, providing a more comprehensive understanding of its downstream regulatory network. Additionally, examining Evi1 function in later developmental stages will be critical to determining its long-term impact on mesodermal derivatives and the formation of the adult body plan. Future research exploring Evi1’s transcriptional targets and downstream effectors will be essential for fully elucidating its role in mesodermal fate decisions and organogenesis.

